# Biophysical characterization of the ETV6 PNT domain polymerization interfaces

**DOI:** 10.1101/2020.08.21.262121

**Authors:** Chloe A. N. Gerak, Sophia Y. Cho, Maxim Kolesnikov, Mark Okon, Michael E. P. Murphy, Richard B. Sessions, Michel Roberge, Lawrence P. McIntosh

**Affiliations:** Department of Biochemistry and Molecular Biology, University of British Columbia, Vancouver, BC, Canada V6T 1Z3; Department of Microbiology and Immunology, University of British Columbia, Vancouver, BC, Canada V6T 1Z3; School of Biochemistry, University of Bristol, Bristol BS8 1TD, United Kingdom; Department of Chemistry, University of British Columbia, Vancouver, BC, Canada V6T 1Z1

**Keywords:** ETS transcription factor family, SAM domain, protein-protein interaction, nuclear magnetic resonance, crystallography, biophysics, molecular dynamics

## Abstract

ETV6 is an ETS family transcriptional repressor that self-associates by its PNT domain to facilitate cooperative DNA binding. Chromosomal translocations frequently generate constitutively active oncoproteins with the ETV6 PNT domain fused to the kinase domain of one of many protein tyrosine kinases. Although an attractive target for therapeutic intervention, the propensity of the ETV6 PNT domain to polymerize via the tight head-to-tail association of two relatively flat interfaces makes it challenging to identify suitable small molecule inhibitors of this protein-protein interaction. Herein we provide a comprehensive biophysical characterization of the ETV6 PNT domain interaction interfaces to aid future drug discovery efforts and help define the mechanisms by which its self-association mediates transcriptional repression. Using NMR spectroscopy, X-ray crystallography, and molecular dynamics simulations, we demonstrate that ETV6 PNT domain variants with monomerizing mutations adopt very stable helical bundle folds that do not change in conformation upon self-association. Amide hydrogen exchange and surface plasmon resonance-monitored alanine scanning mutagenesis studies identified hot spot regions within the self-association interfaces. These regions include both central hydrophobic residues and flanking salt-bridging residues. Collectively, these studies indicate that small molecules targeted to these hydrophobic or charged regions within the relatively rigid interfaces could potentially serve as orthosteric inhibitors of ETV6 PNT domain polymerization.

## Introduction

ETV6 is a modular transcriptional repressor of the ETS (<u>E</u>26 <u>t</u>ransformation <u>s</u>pecific) family for which head-to-tail polymerization of its PNT (or SAM) domain facilitates cooperative binding to tandem DNA sites by its ETS domain (1, 2). The defining DNA-binding ETS domain is conserved among all ETS transcription factors, whereas the PNT domain is present in approximately one-third of ETS paralogs (3). Unlike the monomeric PNT domains of most ETS factors, the ETV6 PNT domain self-associates in a head-to-tail fashion to form an open-ended, left-handed helical polymer (4–6). The only other known self-associating PNT domains in the ETS family are those of *Drosophila* Yan, and possibly human ETV7, as other PNT domains lack suitable interfaces due to amino acid differences or steric blockage (3, 5, 7).

ETV6 is biologically important in embryonic development and hematopoietic regulation (8, 9). Although reported to recruit corepressors such as mSin3A, SMRT, and N-CoR (10), the mechanisms by which ETV6 regulates transcription require further investigation. Polymeric DNA-bound ETV6 is proposed to cause localized chromatin compaction to block access of the transcriptional machinery to target genes. This speculative model is based on the observation that the repeat distance of the helical polymer formed by the PNT domain is comparable to the width of the nucleosome core particle (10).

ETV6 also has preeminent roles in cancer. Frequently, chromosomal translocations fuse gene fragments encoding the PNT domain of ETV6 with the kinase domain of one of many diverse receptor protein tyrosine kinases (PTK) or to the DNA-binding domain from one of several transcription factors (11). Known receptor PTK fusion partners of ETV6 include PDGFβ, JAK2, FGFR3, and NTRK3. The resulting constitutively active self-associated oncoproteins have been linked to over 40 human leukemias, as well as fibrosarcomas, breast carcinomas, and nephromas (11–13).

Due to its presence in numerous fusion oncoproteins, the ETV6 PNT domain is an attractive target for therapeutic intervention (12). However, its propensity to form long insoluble polymers via the tight head-to-tail association (K_D_ ∼ nM) of two relatively flat interfaces hinders identification of suitable small molecule inhibitors (10). These interfaces – termed the ML- and EH-surfaces (mid-loop and end-helix, respectively) – lie roughly on opposite sides on the globular PNT domain. Each is composed of a hydrophobic patch encompassed by polar and charged sidechains. The introduction of an ionizable residue into either hydrophobic patch yields a monomeric PNT domain as judged by several techniques including equilibrium ultracentrifugation and native gel electrophoresis (5, 10). Examples include the V112E or V112R mutations that disrupt the EH-surface or the A93D mutation that disrupts the ML-surface. Two such mutant PNT domains with complementary wild type interfaces can still form a heterodimer. The availability of these monomeric and heterodimeric forms of the PNT domain facilitates studies of ETV6 self-association.

In the case of the ETV6-NTRK3 (EN) fusion oncoprotein, the introduction of monomerizing mutations blocks the ability of EN to polymerize, to activate its PTK, and to transform NIH3T3 cells (14). Furthermore, when co-expressed, the isolated PNT domain has a dominant-negative effect on EN-transformed cells (14). Subsequent studies showed that weakening polymerization by disrupting a peripheral intermolecular salt bridge (K99-D101) also abrogates the ability of EN to transform NIH 3T3 cells (15). Collectively, these studies demonstrate that inhibiting PNT domain polymerization is indeed a viable therapeutic strategy against ETV6-driven cancers.

Understanding the mechanisms of PNT domain polymerization will yield insights in the transcriptional repression properties of ETV6 and the oncogenic properties of ETV6 fusions. In particular, defining structural and thermodynamic differences between monomeric and heterodimeric forms of the PNT domain and determining “hot spot” regions for self-association is important for delineating the mechanisms underlying polymerization and for developing strategies to inhibit this process. Herein, using NMR spectroscopy, X-ray crystallography, and molecular dynamics (MD) simulations, we demonstrated that the structure of the ETV6 PNT domain is very stable and does not change significantly upon self-association. Complementary alanine scanning mutagenesis revealed several hot spot regions within the ML- and EH-surfaces. These residues partake in both hydrophobic and electrostatic interactions and can serve as starting points for targeted rational drug design.

## Results

### PNT domain dimerization characterized by NMR spectroscopy

NMR spectroscopy can give insights into the thermodynamic, kinetic and structural mechanisms of protein-ligand interactions. A particularly convenient approach is to use ^15^N-HSQC spectra to monitor the titration of a ^15^N-labeled protein with an unlabeled, and hence NMR silent, ligand such as another protein (16). Amide chemical shifts are highly sensitive to even subtle environmental changes, and thus an interaction with the unlabeled species can usually be detected through chemical shift perturbations (CSPs) of the labeled protein. Amides exhibiting CSPs typically cluster around the protein-ligand interface, yet may also be distal if binding causes longer range (allosteric) structural changes (16).

To begin this interrogation, purified samples of uniformly ^13^C/^15^N-labeled ETV6 fragments (residues 40-125) that contain either an A93D mutation or V112E mutation (henceforth described as either the A93D- or V112E-PNT domains) were prepared for NMR spectroscopic characterization. Under neutral pH solution conditions, the A93D- and V112E-PNT domains yielded well-dispersed NMR spectra indicative of stably folded structures (Figs. 1A and 2A). However, the latter showed some propensity to self-associate, and improved spectra were obtained at a sample pH value of 8.0. Presumably, this reflects the deprotonation of E112, which may have an anomalously high p*K*_a_ value when buried at the polymer interface. Unfortunately, the more alkaline conditions resulted in some loss of signal intensity due to base-catalyzed amide hydrogen exchange (HX).

**Figure 1.**
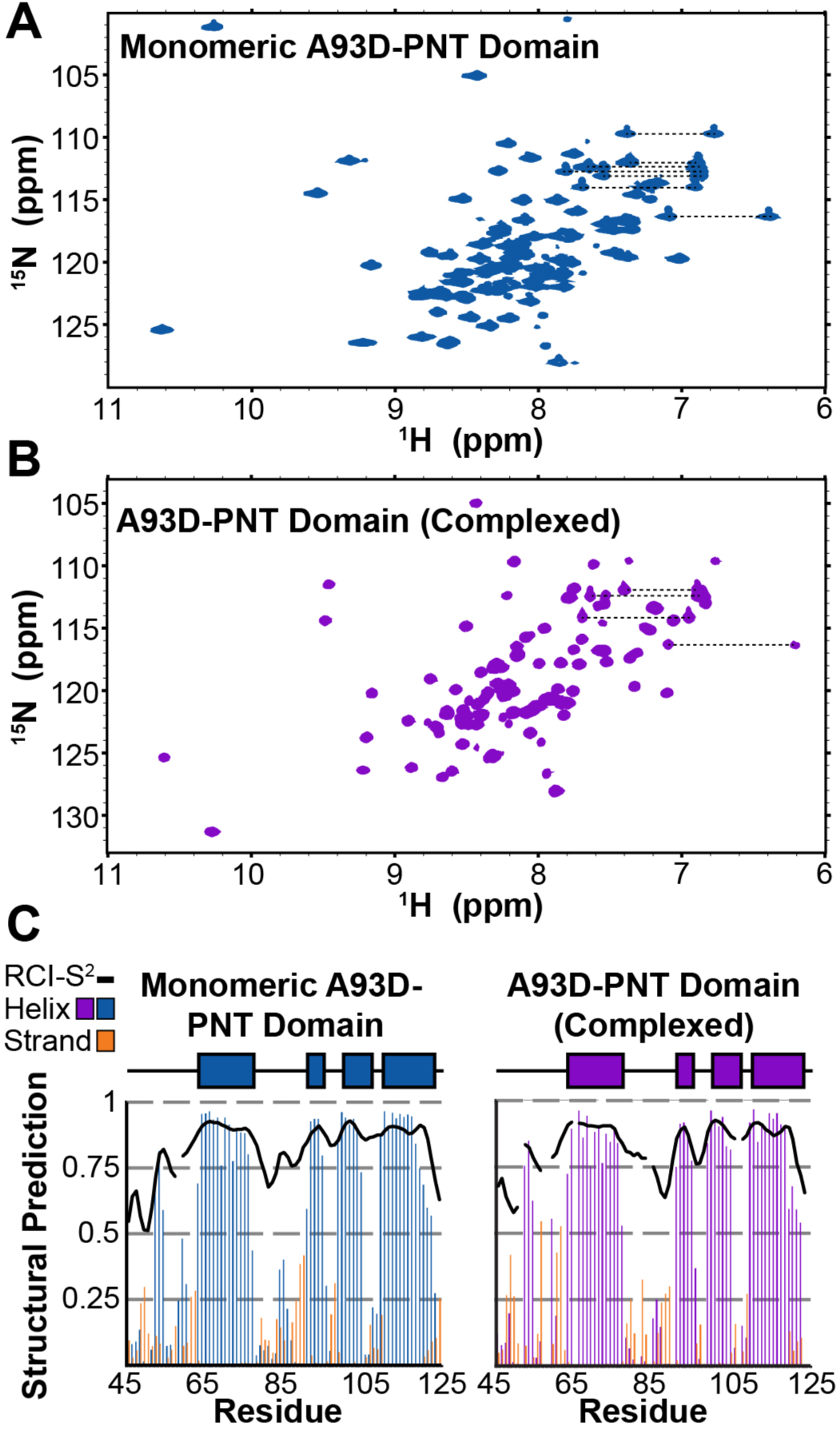
NMR spectra of the A93D PNT domain in its monomeric and complexed states. The ^15^N-HSQC spectra of the ^15^N-labeled A93D-PNT domain in it monomeric form **(A)**, and as a heterodimer with the unlabeled V112E-PNT domain **(B)**, recorded in 20 mM MOPS, 50 mM NaCl, 0.5 mM EDTA and 5% D_2_O at pH 7.0 and 25 °C. Assignments are provided in Figs. S2 and S4, respectively. **(C)** The predicted secondary structural elements and RCI-S^2^ values (black lines; decreasing values from 1 to 0 indicate increasing flexibility) for the A93D-PNT domain in its monomeric (left) and complexed (right) states, calculated with the MICS algorithm. Missing data correspond to residues lacking chemical shift assignments. The locations of the four α-helices (H1-H4) observed in the X-ray crystal structures of the self-associated ETV6 PNT domain are indicated as rectangles above each plot.

Upon addition of the unlabeled V112E PNT domain to the ^15^N-labeled A93D-PNT domain, many amides exhibited CSPs in the slow exchange regime (Fig. 1B). That is, at an intermediate titration point, separate ^1^H^N^-^15^N peaks corresponding to the unbound (monomeric) and bound (heterodimeric) forms of the labeled protein were observed. This is consistent with the previously reported K_D_ value ∼ 2 nM for the high-affinity binding equilibrium (10). Comparable results were observed for the reciprocal titration of the unlabeled A93D-PNT domain to the ^15^N-labeled V112E-PNT domain (Fig. 2A and Fig. S5). Parenthetically, although the oligomerization states of the PNT domains in these experiments were not directly determined, the results are entirely consistent with previous studies showing that the A93D- and V112E-PNT domains are monomeric when separated and heterodimeric when combined (10).

**Figure 2.**
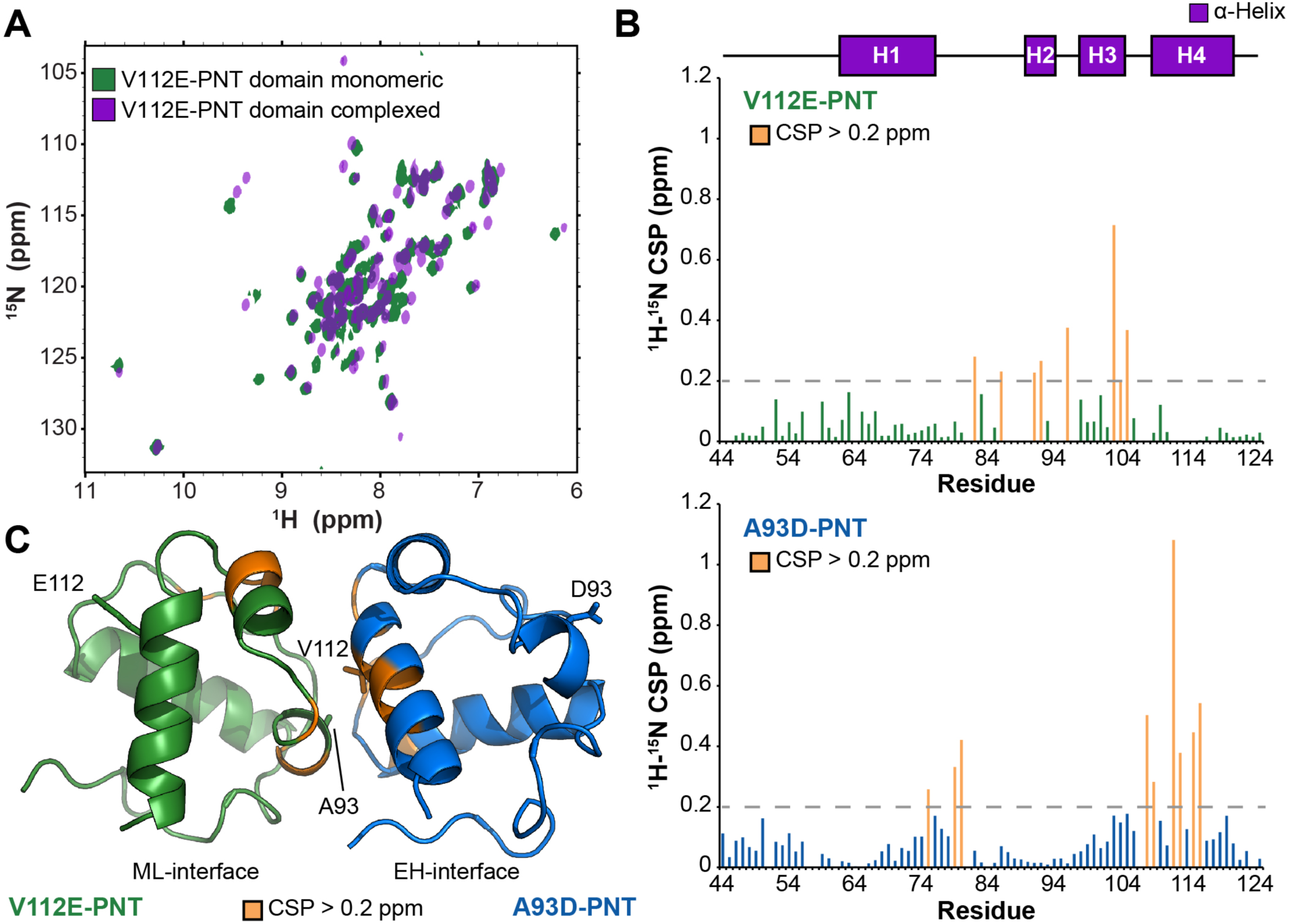
PNT domain dimerization interface identified by amide chemical shift perturbations. **(A)** Overlaid ^15^N-HSQC spectra of the ^15^N-labeled V112E-PNT domain in the absence (green) and presence of a 1.1 molar excess of the unlabeled A93D-PNT-domain (purple). Assignments are provided in Figs. S1 and S3, respectively. **(B)** Backbone amide ^1^H^N^-^15^N CSPs resulting from the heterodimerization of the V112E-PNT (top) and A93D-PNT (bottom) domains. Missing data correspond to prolines or residues without assigned NMR signals in either protein state. Most residues showed small CSPs which may be due in part to the different conditions under which spectra were assigned (V112E-PNT domain monomer, pH 8.0; V112E-PNT domain complexed with the A93D-PNT domain, pH 7.5; A93D-PNT domain monomer, pH 7.0; A93D-PNT domain complexed with the V112E-PNT domain, pH 7.5). **(C)** However, amides with CSP values > 0.2 ppm, which are highlighted in orange on a model of the V112E- (green) / A93D- (blue) PNT domain heterodimer, map to the interfacial regions. The cartoon is derived from PDB: 1LKY with a V112R-PNT domain.

Through a combination of scalar and NOE correlation experiments, NMR signals were assigned from most main chain ^1^H, ^13^C, and ^15^N nuclei of the V112E- and A93D-PNT domains in their monomeric forms and as heterodimers with their unlabeled partners (Figs. S1-S4). Utilizing the MICS algorithm (17), the secondary structural elements for the four species were predicted from these chemical shifts. In each case, four distinct helical regions were detected (Fig. 1C and Fig. S6). These coincide well with the four α-helices (H1: R63-E76; H2: G91-L94; H3: K99-R105; H4: G110-K122) identified in the X-ray crystal structures of the self-associated ETV6 PNT domains (PDB: 1JI7 and 1LKY) by PDBsum (18). Matching two short N-terminal 3_10_-helices observed in these crystal structures, residues A52-L54 and, to a lesser extent, P58-Y60 also have chemical shifts indicative of helical character. In contrast, such diagnostic chemical shifts were not seen for residues S84-T86 even though they are classified as a forming a 3_10_-helix in a subset of the monomer subunits of PDB file 1LKY. This minor discrepancy may arise because these residues are within an extended, solvent exposed polypeptide segment between helices H1 and H2. Amides in this region have chemical shift-derived random coil index-squared ordered values (RCI-S^2^, a proxy for backbone dynamics (19)) indicative of increased flexibility relative to the well-ordered helices (Fig. 1C and Fig. S6). Most importantly, these analyses demonstrated that the A93D- and V112E-PNT domains have very similar secondary structures in their monomeric and heterodimeric forms. Thus, the proteins in solution do not undergo any significant changes in secondary structure upon mutation or head-to-tail association into the structures previously characterized by X-ray crystallography.

Armed with chemical shift assignments, the amide ^1^H^N^-^15^N CSPs resulting from PNT domain dimerization were readily calculated. Of note, residues in the ^15^N-labeled A93D- and V112E-PNT domains that experienced the greatest CSP cluster within the EH- and ML-surfaces, respectively (Figs. 2B and 2C). This confirms that, as seen by X-ray crystallography, the two monomerized PNT domains indeed associate in solution through their wild type interfaces.

### Crystallographic comparison of monomeric and dimeric PNT domains

In their original studies, the Bowie group obtained crystals of the V112E-PNT domain (PDB: 1JI7, *C*2 space group, 3 monomers in the asymmetric unit) (10). Despite burial of E112, the monomers assembled in the crystal lattice via their ML- and EH-surfaces to form an extended helical polymer with an approximate 6_5_ screw symmetry (Fig. 3A). Subsequently, they determined the structure of a heterodimer composed of a A93D-PNT domain bound to a V112R-PNT domain via their complementary wild type interfaces (PDB: 1LKY, *P*1 space group, 3 heterodimers in the asymmetric unit) (6). Although no longer polymeric within the crystal lattice, a model built from PDB: 1LKY using appropriate monomers subunits with native interfaces closely matched the polymeric structure of PDB: 1JI7 with the V112E substitution. Thus, the latter serves as a reliable experimental structure of the ETV6 PNT domain polymer.

**Figure 3.**
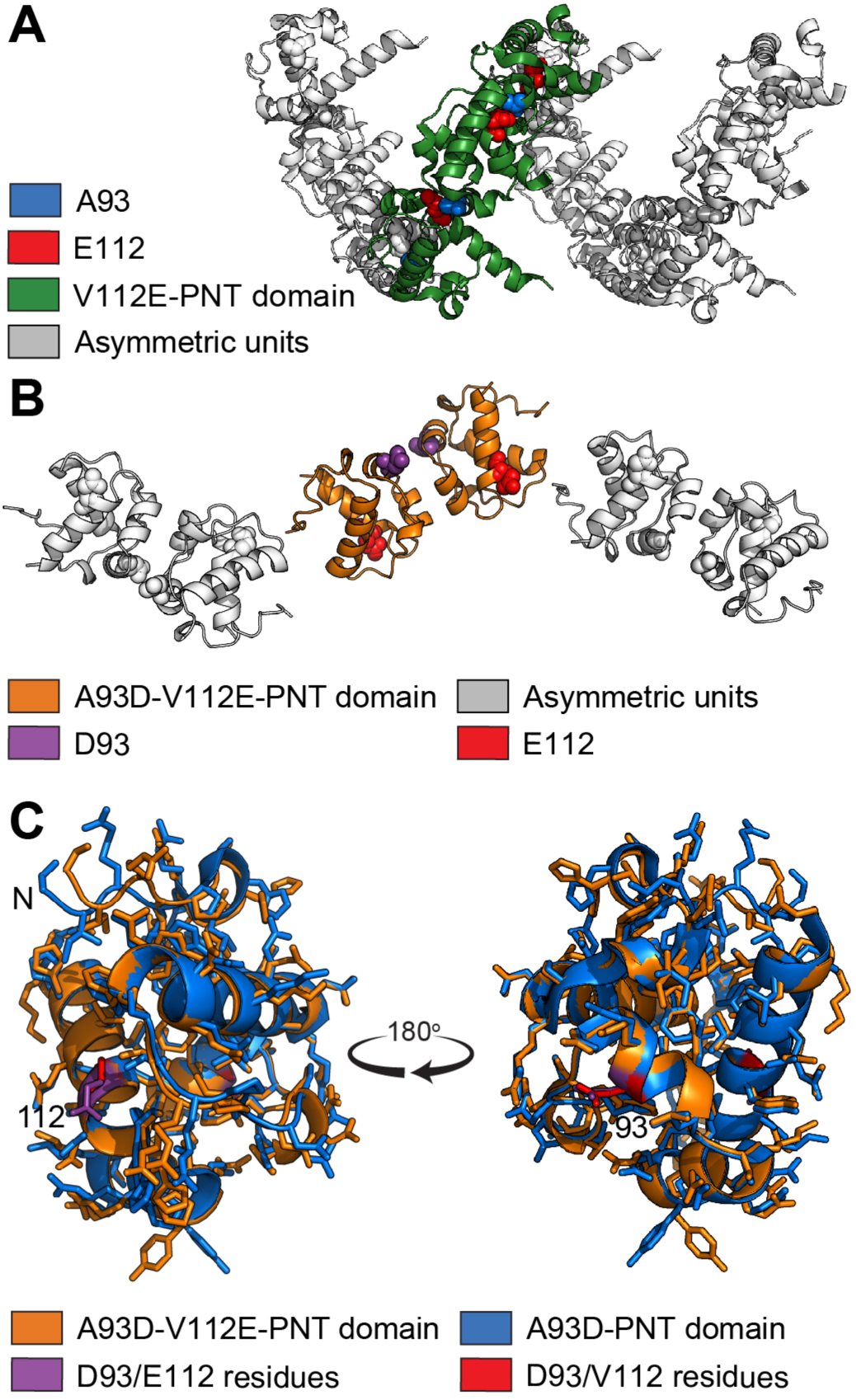
Structural comparison of the monomeric and polymeric ETV6 PNT domain variants. **(A)** The X-ray crystallographic structure of the V112E-PNT domain from PDB: 1JI7. The three subunits of the V112E-PNT domain in the asymmetric unit (green) form a helical polymer with neighboring subunits (grey). The mutated E112 sidechain is shown as red spheres and the wild type A93 as blue spheres. (B) The structure of the A93D-V112E-PNT domain determined herein. The two A93D-V112E-PNT domains in the asymmetric unit are highlighted in orange (E112 as red spheres, D93 as purple spheres), and neighboring asymmetric units are in grey. The crystal contacts are unlike those in the polymeric structure and no wild type protein-protein polymerization interface is seen. **(C)** Structural overlay of an A93D-PNT domain subunit that was in complex with a V112R-PNT domain (blue, PDB: 1LKY chain B) and the A93D-V112E-PNT domain (orange), determined herein. Residues at positions 93 and 112 are highlighted. In contrast to well aligned interior side chains, variations in the rotamer conformations of surface sidechains (e.g. the lower tyrosine) and the N-terminal residues can be seen.

The NMR spectroscopic studies presented above indicate that the secondary structures and binding interfaces of the A93D- and V112E-PNT domains in solution closely resemble those observed by X-ray crystallography. However, the PNT domain in a complexed form may have subtle structural differences relative to a monomeric form. To examine this possibility, a PNT domain with both A93D and V112E mutations was generated and found to crystallize in 2.8 M sodium acetate at pH 7.0. Its structure was solved to 1.86 Å resolution using molecular replacement (Table S2). The A93D-V112E-PNT domain crystallized in the space group *P*6_5_22 with two monomers in the asymmetric unit (Fig. 3B). More importantly, the presence of both monomerizing mutations prevented any intermonomer interactions within the crystal lattice via the MH- and EL-surfaces. Rather, nearest neighbor contacts were via alternative interfaces that are not functionally relevant.

Regardless of differing crystallization conditions and mutations, the structure of the A93D-V112E-PNT domain closely resembles those previously determined for the ETV6 PNT domain. Using the DALI server to compare one subunit from this structure with a A93D-PNT domain subunit from PBD entry 1LKY, a total of 77 residues were aligned with a RMSD value of 0.7 Å and Z-score of 16.8 (20). However, a few subtle differences can be seen upon detailed comparison (Fig. 3C). For example, residues N-terminal to helix H1 have variable conformations. This is consistent with their RCI-S^2^ scores indicating a degree of flexibility (Fig. 1C and Fig. S6). Not unexpectedly, several surface residues adopted different sidechain rotamer conformations, whereas residues within the interior hydrophobic core of the PNT domain superimposed well. Most importantly, the local structural features of the ML (including residue 93) and EL (including residue 112) interfaces do not differ despite the presence or absence of monomerizing mutations or their association upon dimer or polymer formation. Thus, interactions of the ETV6 PNT domain do not contribute to any discernible conformational changes between its monomeric, heterodimeric, or polymeric forms.

### Amide hydrogen exchange data show increased protection of interfacial residues upon dimerization

Amide HX is a useful technique to characterize protein structure, stability and dynamics, as well as identifying ligand-binding interfaces (21–23). Through a continuum of local to global conformational fluctuations, main chain amide hydrogens are constantly exchanging with the hydrogens of solvent water (24). If a labile amide proton exchanges for a deuteron it will become silent for ^1^H-detected NMR, and thus its signal will disappear from an ^15^N-HSQC spectrum. The rate at which it disappears is determined by its structural features (e.g. hydrogen bonding and solvent accessibility) as well as the experimental conditions (e.g. pH and temperature). To account for the latter, the observed exchange rate constant can be compared to the predicted rate constant for a random coil polypeptide with the same sequence and under the same conditions. The ratio of the predicted versus observed rate constants is the protection factor (PF).

To gain further insights into the ETV6 PNT domain dynamics, NMR spectroscopy was used to measure the amide PFs of its monomeric and heterodimeric forms. By comparing the spectra recorded after three days of exchange (Fig. 4A), it is immediately obvious that substantially more amides were protected from HX in the heterodimeric versus monomeric species. Exchange rate constants for most amides were determined by fitting their time dependent ^1^H^N^-^15^N signal intensities to single exponential decays. These values were converted into the PFs shown in Figs. 4B and 4C (and Table S3).

**Figure 4.**
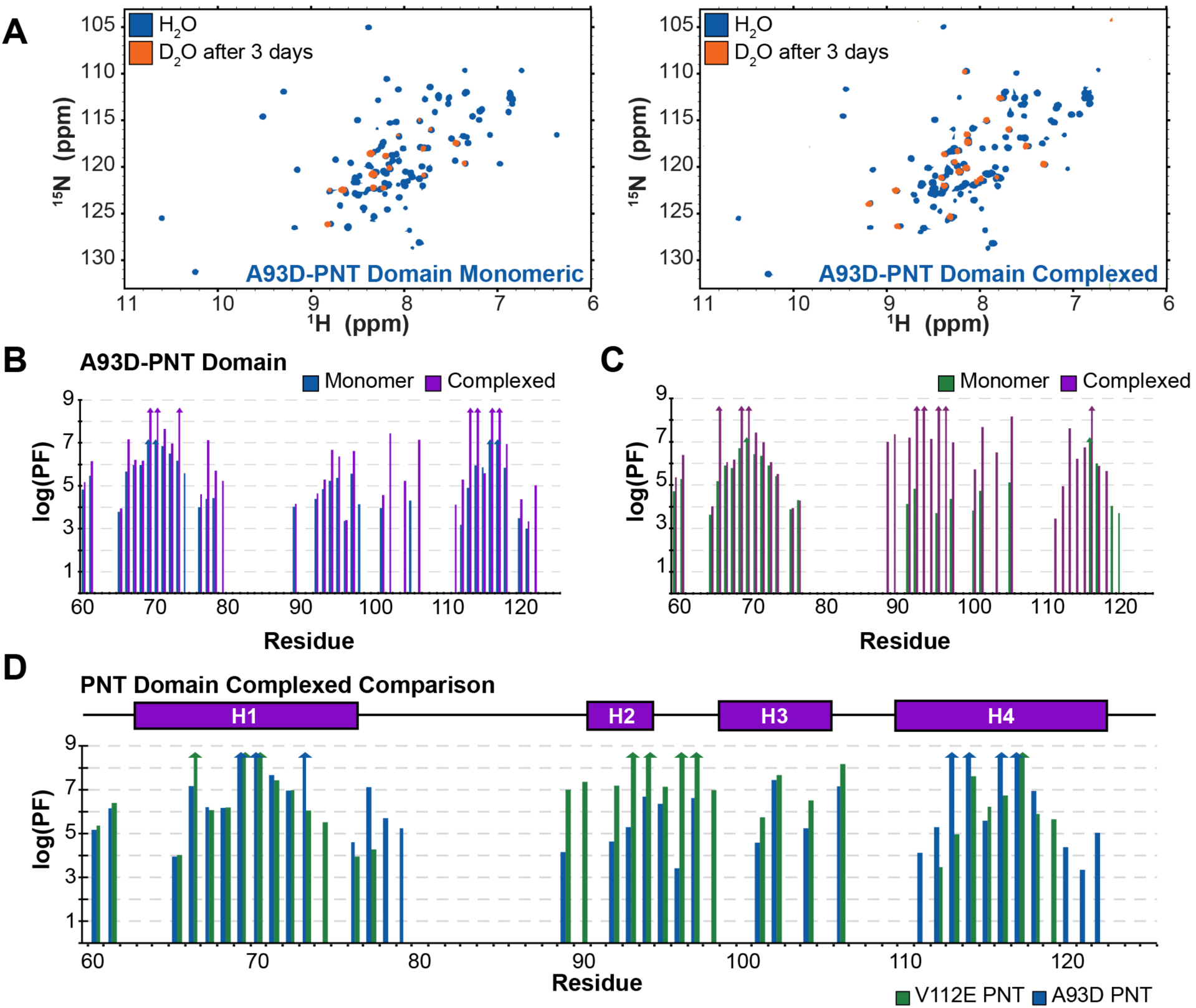
PNT domain amide HX protection factors increase upon heterodimerization. **(A)** Overlaid ^15^N-HSQC spectra for the monomeric (left) and complexed (right) forms of the A93D-PNT domain in H_2_O (blue) and after 3 days at 21 °C in D_2_O buffer (orange). Significantly more amides in the heterodimeric species were protected from HX. A summary of the PFs for the A93D-PNT **(B)** and V112E-PNT **(C)** domain in its monomeric and complexed states. Missing data correspond to prolines, residues with unassigned or overlapping amide chemical shifts, or residues that exchanged prior to collection of the first ^15^N-HSQC spectrum after transfer into D_2_O buffer and thus have log(PF) values less than an estimated upper limit of ∼ 3. The latter include amides preceding residue 60, which were not observed for any protein after transfer into D_2_O buffer. In the cases of amides with that had not exchanged significantly after 3 months, estimated lower limits of log(PF) values > 7 (monomeric) and > 8.5 (heterodimeric) are indicated by upwards arrows. **(D)** A comparison of the amide PFs of the V112E PNT-domain and A93D-PNT domains when associated within a heterodimer. The most protected amides are located near the complementary ML- (containing A93) and EH- (containing V112) interfaces of the two species, respectively. The helical regions are indicated with rectangles.

The monomeric A93D- and V112E-PNT domains exhibited very similar patterns of amide HX. All residues N-terminal to Y60 and most of those in interhelical regions exchanged rapidly under these experimental conditions. This is consistent with their surface exposure and general lack of intramolecular hydrogen bonding interactions. Conformational flexibility of these residues was also indicated by their RCI-S^2^ values (Fig. 1C and Fig. S6). Conversely, amides within or near the four α-helices showed substantial protection from HX. In particular, W69 and L70 in helix H1 and L116 and L117 in helix H4 of both proteins exchanged very slowly, with several of these residues having log(PF) values > 7. Under the commonly observed EX2 conditions, with pH-dependent bimolecular kinetics, PFs reflect the residue-specific free energy changes, ΔG_HX_ = 2.303RTlog(PF), governing local or global conformational equilibria leading to exchange (25). Assuming that these most protected amides exchange through global or near-global structural fluctuations, these HX data provide an estimation of the unfolding free energy for each monomeric PNT domain of > 40 kJ/mol. Such a value is consistent with the view that, even without polymerizing, the ETV6 PNT domain adopts a very stable folded conformation. This further indicates that the A93D and V112E mutations prevent polymerization without disrupting the structure or stability of the monomeric PNT domain.

Heterodimerization resulted in increased HX protection for many residues in both the A93D- and V112E-PNT domains (Figs. 4B and 4C). Indeed, several amides in both proteins did not exchange significantly even after 3 months in D_2_O buffer and thus have log(PF) > 8.5 (and ΔG_HX_ > 48 kJ/mol). Although global stabilization upon heterodimer formation is expected, many of the residues with at least a 100-fold increase in HX protection localized around the complementary interfacial regions of the two PNT domains (Figs. 4D, 5A and 5B). These are exemplified by amides within or near the ML-surface of the V112E-PNT domain (N90, K92, A93, L96, T98, D101, F102) and the EH-surface of the A93D-PNT domain (F77, V112, L113, Y114). Given that the structures of the A93D- and V112E-PNT domains do not change significantly upon heterodimerization, the increased protection of interfacial residues against HX may result from their local stabilization against conformational fluctuations allowing exchange. Alternatively, if exchange occurs predominantly through transient monomers, then the increased protection of amides in the heterodimer would reflect the equilibrium population distribution of these two species (21). Regardless of mechanism, the HX data are consistent with the role of these interfaces in ETV6 PNT domain polymerization.

**Figure 5.**
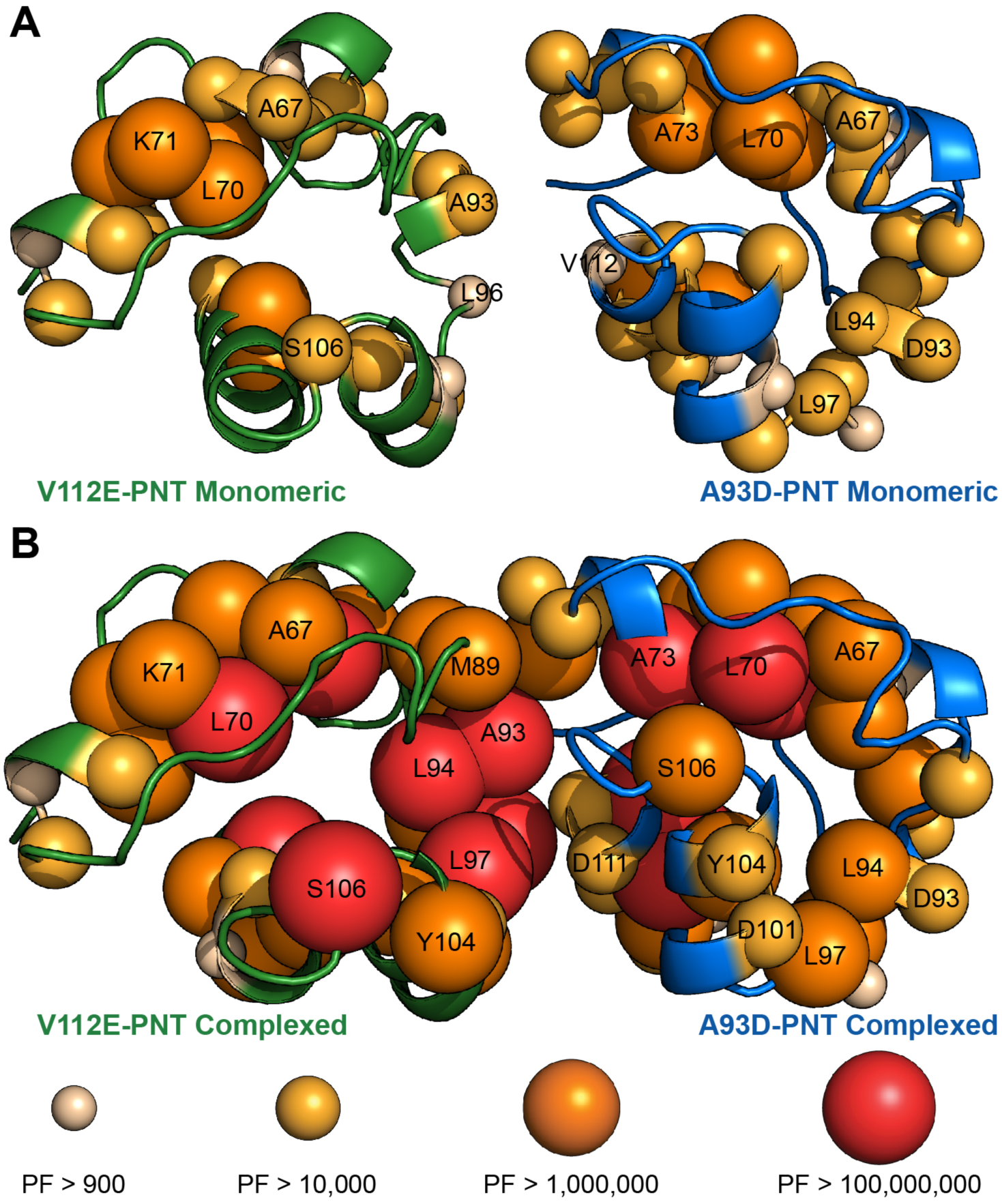
Mapping of HX protection factors on the monomeric and complexed PNT domain structures. Amide PFs are displayed as spheres on the ribbon diagrams (from PDB: 1LKY) of the V112E- and A93D-PNT domains in their (**A**) monomeric and (**B**) heterodimeric forms according to the indicated color and size scheme. Residues without spheres are either prolines, lack an assigned or fully resolved ^1^H^N^- ^15^N signal, or exchanged too fast for reliable HX quantitation. Both monomeric PNT domains showed similar HX profiles, with the most protected amides located in structured helical regions. Upon association, the PFs of many amides at the complementary dimer interfaces increased.

### Alanine scanning mutagenesis at the PNT domain self-association interface

Alanine scanning mutagenesis was used to identify which residues at the PNT domain heterodimer interface contribute most to binding affinity. The mutation of a residue to alanine reduces its side chain to a single methyl group, thereby eliminating its contributions to intermolecular binding, while also avoiding the introduction of any additional non-native interactions. Alanine was chosen over glycine as the latter may lead to increased backbone flexibility. For these studies, SPR was used to characterize the binding interaction as this technique is rapid and uses only small quantities of bacterially expressed proteins.

Initial controls were performed to reproduce the results of previously reported SPR studies of the PNT domain dimerization (10). Either the biotinylated A93D- or V112E-PNT domain was immobilized on a streptavidin chip, and the complementary (positive control) or same (negative control) PNT domain was applied as the analyte at various concentrations (Fig. S7). High affinity binding was seen for the heterodimer pairings, whereas the identical PNT domains did not measurably interact. The A93D-PNT domain analyte bound the V112E-PNT domain ligand with a fit K_D_ value of 7.5 nM, and the V112E-PNT domain analyte bound the A93D-PNT domain ligand with a fit K_D_ value of 5.1 nM. These results agreed well with previously reported K_D_ values of 1.7 to 4.4 nM for the PNT domain interactions as measured by SPR (10) and ITC (15). Thus, SPR was used to characterize the effects of alanine substitutions of 18 residues within or around the EH-surface of the A93D-PNT domain and 14 residues within or around the ML-surface of the V112E-PNT domain (Tables 1 and 2).

**Table 1.** Alanine scanning mutagenesis of the EH-surface on the A93D-PNT domain ^*a*^.

| **Ala Mutation** | **$k_{on}$ ($M^{-1}s^{-1}$)** | **$k_{off}$ ($s^{-1}$)** | **$K_D$ (nM)** | **$\Delta\Delta G$ (kJ/mol) <sup>b</sup>** |
| --- | --- | --- | --- | --- |
| None | $2.0 \times 10^5$ | $1.5 \times 10^{-3}$ | 7.5 | - |
| S47A | $2.1 \times 10^5$ | $1.4 \times 10^{-3}$ | 6.9 | -0.2 |
| I48A | $3.4 \times 10^5$ | $4.0 \times 10^{-3}$ | 12 | 1.2 |
| E76A | $6.2 \times 10^5$ | $6.2 \times 10^{-3}$ | 9.9 | 0.7 |
| F77A | $3.4 \times 10^4$ | $1.5 \times 10^{-2}$ | 450 | 10.2 |
| S78A | $2.8 \times 10^6$ | $9.3 \times 10^{-3}$ | 3.3 | -2.0 |
| L79A | $8.1 \times 10^3$ | $1.6 \times 10^{-3}$ | 200 | 8.2 |
| R80A | $9.1 \times 10^4$ | $2.9 \times 10^{-3}$ | 32 | 3.6 |
| K99A | $2.4 \times 10^3$ | $2.3 \times 10^{-3}$ | 930 | 12.0 |
| E100A | $3.1 \times 10^5$ | $1.5 \times 10^{-3}$ | 5 | -1.0 |
| R103A | $2.5 \times 10^5$ | $5.4 \times 10^{-3}$ | 21 | 2.6 |
| P107A | $2.6 \times 10^5$ | $2.1 \times 10^{-3}$ | 8.1 | 0.2 |
| H108A | $2.6 \times 10^5$ | $1.4 \times 10^{-3}$ | 5.4 | -0.8 |
| D111A | $3.3 \times 10^3$ | $1.4 \times 10^{-3}$ | 400 | 9.9 |
| V112A | $2.0 \times 10^4$ | $1.1 \times 10^{-2}$ | 550 | 10.7 |
| Y114A | $3.5 \times 10^4$ | $4.5 \times 10^{-3}$ | 130 | 7.1 |
| E115A | $2.3 \times 10^5$ | $2.5 \times 10^{-3}$ | 11 | 1.0 |
| L116A | $2.3 \times 10^5$ | $7.6 \times 10^{-4}$ | 3.3 | -2.0 |
| H119A | $2.7 \times 10^5$ | $3.4 \times 10^{-3}$ | 13 | 1.4 |

<sup>a</sup> All A93D-PNT domain analytes were run on the streptavidin SPR chip bound with the biotinylated V112E-PNT domain ligand.
<sup>b</sup> Calculated as $\Delta\Delta G = RT \ln(K_{D,mutant}/K_{D,wild\ type})$ where wild type is the top-listed protein with an unmodified interface.

**Table 2.** Alanine scanning mutagenesis of the ML-surface on the V112E-PNT domain ^*a*^.

| **Ala Mutation** | **$k_{on}$ ($M^{-1} s^{-1}$)** | **$k_{off}$ ($s^{-1}$)** | **$K_D$ (nM)** | **$\Delta\Delta G$ (kJ/mol) <sup>b</sup>** |
| --- | --- | --- | --- | --- |
| None | $4.4 \times 10^5$ | $2.3 \times 10^{-3}$ | 5.1 | - |
| I59A | $4.3 \times 10^5$ | $2.5 \times 10^{-3}$ | 5.8 | 0.3 |
| R63A | $3.6 \times 10^5$ | $5.4 \times 10^{-3}$ | 15 | 2.7 |
| N85A | $3.5 \times 10^5$ | $3.4 \times 10^{-3}$ | 9.9 | 1.6 |
| E88A | $5.0 \times 10^5$ | $4.0 \times 10^{-3}$ | 8.1 | 1.1 |
| M89A | $9.7 \times 10^4$ | $3.3 \times 10^{-2}$ | 340 | 10.4 |
| N90A | $7.8 \times 10^4$ | $3.7 \times 10^{-2}$ | 480 | 11.3 |
| K92A | $3.5 \times 10^5$ | $2.2 \times 10^{-2}$ | 62 | 6.2 |
| L96A | $9.5 \times 10^3$ | $9.5 \times 10^{-3}$ | 1000 | 13.1 |
| L97A | $9.8 \times 10^4$ | $6.8 \times 10^{-2}$ | 700 | 12.2 |
| T98A | $6.2 \times 10^5$ | $5.4 \times 10^{-3}$ | 8.8 | 1.3 |
| E100A | $5.0 \times 10^5$ | $3.5 \times 10^{-3}$ | 7 | 0.8 |
| D101A | $6.3 \times 10^3$ | $2.5 \times 10^{-3}$ | 400 | 10.8 |
| Y104A | $7.3 \times 10^5$ | $1.5 \times 10^{-2}$ | 21 | 3.5 |
| R105A | $2.3 \times 10^3$ | $5.4 \times 10^{-3}$ | 2340 | 15.2 |

<sup>a</sup> All V112E-PNT domain analytes were run on the streptavidin SPR chip bound with the biotinylated A93D-PNT domain ligand.
<sup>b</sup> Calculated as $\Delta\Delta G = RT \ln(K_{D,mutant}/K_{D,wild\ type})$ where wild type is the top-listed protein with an unmodified interface.

Six out of 18 on the A93D-PNT domain and 7 out of 14 tested residues on the V112E-PNT domain had large detrimental effects on binding (ΔΔG > 6 kJ/mol) and can be classified as hot spots (Fig. 6A). Although more difficult to interpret than K_D_ values, these mutations acted to varying degrees by slowing the association rate constants, k_on_, and/or increasing the dissociation rate constants, k_off_. The higher proportion of hot spot interfacial residues on the V112E-PNT domain, including the three most detrimental of the entire set of alanine mutants, suggests that the features of this interface contribute most significantly to PNT domain polymerization.

**Figure 6.**
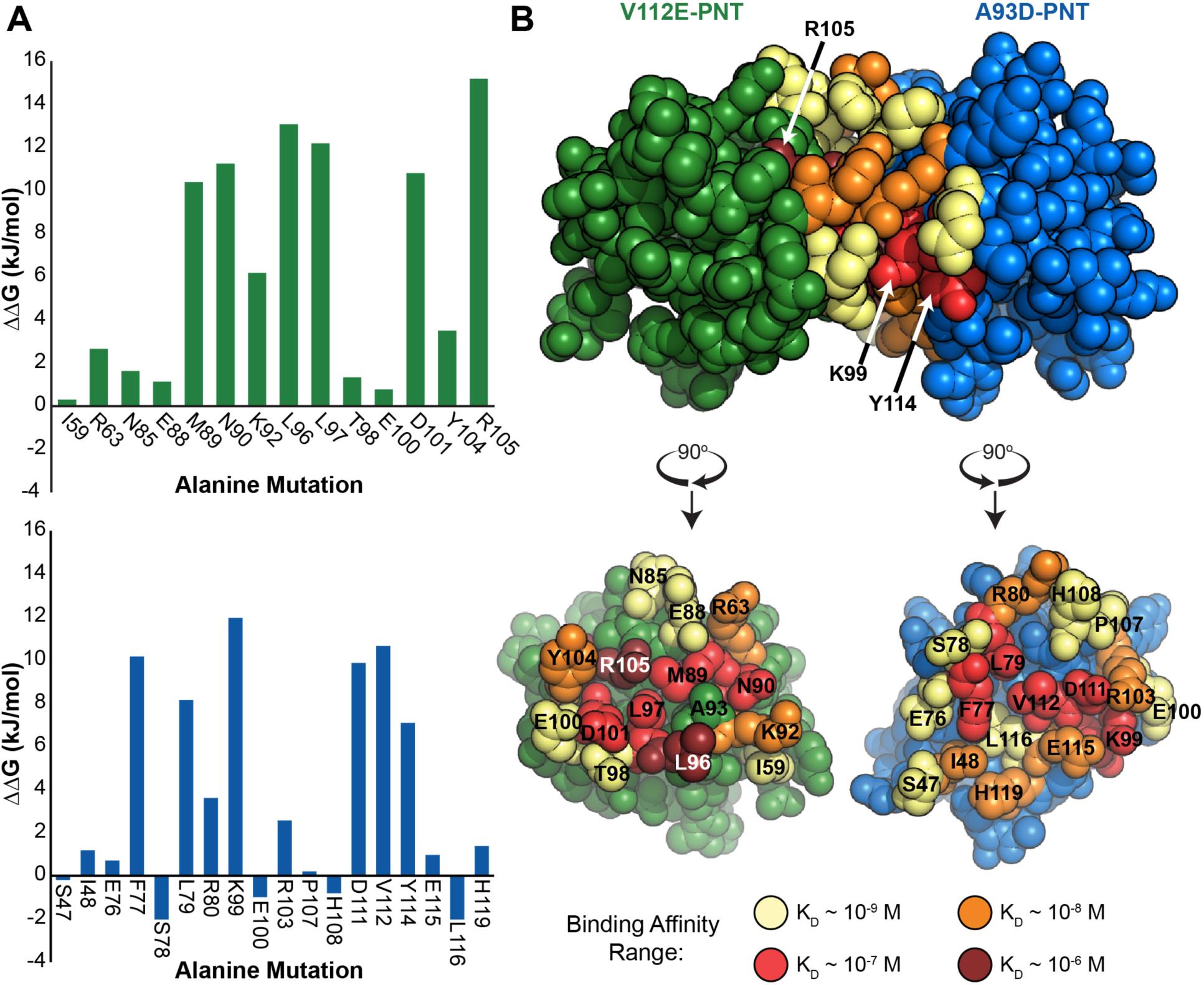
Characterization of the PNT domain interface by alanine scanning mutagenesis. **(A)** A summary of the effects of alanine substitutions on the ML-surface of V112E-PNT domain (top) and the EH-surface of the A93D-PNT domain (bottom) interfaces. Each ΔΔG value corresponds to the change in binding free energy to the complementary PNT domain relative to that of the reference protein with an unmodified wild type interface (Tables 1 and 2). **(B)** The effects of alanine substitutions on the K_D_ values for heterodimer dissociation are mapped onto the structures of the V112E- (green background) and A93D- (blue background) PNT domains. Residues for which alanine mutations maintained binding affinity approximately the wild type K_D_ (∼ 10^−9^ M) are indicated in pale yellow, whereas those weakening binding by approximately 10-fold (orange), 100-fold (red) and 1000-fold (burgundy) are in increasingly darker colors.

Many of the hydrophobic residues at the center of the heterodimer interfaces are hot spots for binding. In particular, the L96A mutation on the ML-surface of the V112E-PNT domain resulted in an ∼1000-fold decrease in affinity. As shown in Fig. 6B, L96 protrudes out of the V112E-PNT domain as a “bump” to fit into the “hole” between residues F77, V112, E115 and H119 on the EH-surface of the A93D-PNT domain. Thus, in addition to contributing to heterodimerization through the hydrophobic effect, L96 partakes in favorable van der Waals interactions at the protein-protein interface. Other hydrophobic residues that are important include L97 and M89 on the ML-surface and F77, L79, and V112 on the EH-surface.

Surrounding the hydrophobic center of the heterodimer interface is a ring of residues that generally have reduced contributions towards binding affinity (Fig. 6B). However, an important exception is the peripheral K99 on the A93D-PNT domain EH-surface. This residue forms an intermolecular salt bridge with D101 on the ML-surface of the V112E-PNT domain. Alanine mutations of K99 or D101 resulted in decreased binding affinities of 930 nM or 400 nM, respectively. Thus, consistent with their salt bridge pairing, either mutation weakened binding relative to the wild type by a factor of ∼ 100. Mutation of K99 has been shown to interfere with EN oncogenic cellular transformation, confirming the functional importance of this salt bridge (15). A second intermolecular salt bridge consistently seen in the PNT domain heterodimer structures involves R105-D111. The R105A and D111A mutations also severely weakened binding to K_D_ values of 2.3 μM and 400 nM, respectively, thus confirming the importance of this interaction. In contrast, several additional salt bridges involving K99-E100, R103-E100, and R103-D101 are seen in some, but not all crystallographically defined interfaces. However, alanine substitutions of either E100 or R103 did not significantly impact binding. Thus, the K99-D101 and R105-D111 salt bridges play important roles in PNT domain association, whereas those involving E100 or R103 do not. It is also notable that the polar residue N90 on the ML-surface of the V112E-PNT domain surface is also a hot spot, but it does not appear to have a potential reciprocal hydrogen bond donor or acceptor. Overall, the alanine scanning mutagenesis study illustrated that PNT domain dimerization is a result of interactions involving both hydrophobic and charged interfacial residues.

### Molecular dynamics simulations of the PNT monomer and dimer

MD simulations can yield insights into the conformational flexibility and motions of a protein. Therefore, simulations of 100 ns were run on the PNT domain heterodimer (an A93D-PNT domain and V112R-PNT domain subunit from 1LKY.pdb) and the A93D-V112E-PNT domain monomer, reported herein. Over this time period, both the heterodimer and the monomer remained structurally stable with root mean square deviation (RMSD) ∼ 2 Å for all non-hydrogen atoms (Fig. 7A). This is typical for globular proteins of this size. Importantly, the PNT domain subunits remained associated with each other, indicating that the heterodimer can be modelled *in silico*. Comparing the averaged root mean square fluctuation (RMSF) of each residue throughout the simulation also showed little flexibility except at the termini (Fig. 7B). With the A93D-PNT domain, there was a slight spike in the RMSF of residue E88 and this likely due to the residue being central on a loop region. Aside from these minor exceptions, there appeared to be no correlation of the RMSF of a residue and whether it is present in a helix or loop region. Collectively, these MD simulations indicated that both the PNT domain monomers and heterodimer are structurally stable and do not experience any major conformational fluctuations on the sub-microsecond timescale.

**Figure 7.**
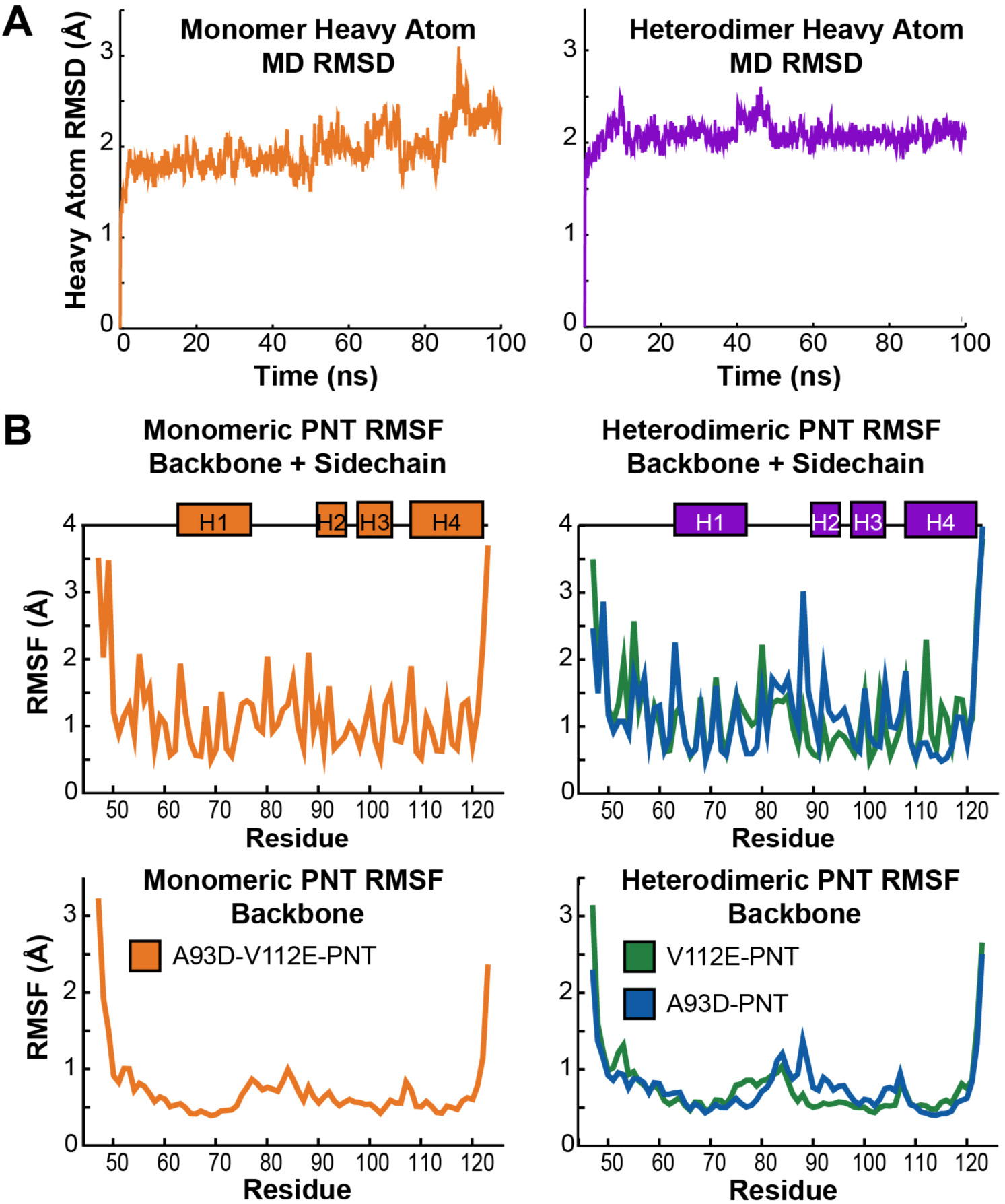
Analysis of PNT domain structural fluctuations from 100 ns MD simulations. **(A)** The RMSD values of non-hydrogen atoms throughout the 100 ns MD simulation are plotted relative to the starting structures. After the initial energy minimization, both the monomer (A93D-V112E-PNT domain) and heterodimer complex showed little conformational deviations. **(B)** Plotted are the RMSF values for non-hydrogen atoms in each residue (top, backbone and sidechain; bottom, backbone only) throughout the 100 ns MD simulation. The values are relative to the average side chain conformations after superimposition of the helical regions of the PNT domain. The left plots show the A93D-V112E-PNT monomer (orange) and the right plots show the heterodimer of the V112E-PNT domain (green) and A93D-PNT domain (blue). With the exception of terminal residues, there are very little fluctuations throughout the simulation.

Principal component (PC) analysis of the trajectories was also performed to explore the low frequency conformational motions of the monomeric and heterodimeric dimeric PNT domains. The proportion of the motion (variance) is largely confined to the first few principal components (Fig. S8A). Inspection of the first two principal components (see Supplemental Movies 1 to 4) shows that the motion in each case mostly comprises random conformational changes of the N- and C-termini. Projecting the trajectories onto the principal components (Fig. S8B) indicates that many combinations of PC1 and 2 are accessed and thus the MD simulations provided a good sampling of the conformational space available to the terminal residues of the ETV6 PNT domain.

## Discussion

### The ETV6 PNT domain retains a similar structure in monomeric and heterodimeric states

These biophysical studies of the PNT domain show that its structure is very stable in the monomeric and heterodimeric forms, and is not dependent upon the presence of monomerizing mutations. There are no substantial conformational differences between the species as seen both by X-ray crystallography and by NMR spectroscopy (secondary structural propensities and chemical shift perturbations). Furthermore, the X-ray crystallographic structure of a monomeric A93D-V112E-PNT domain closely resembles that of the heterodimers, and does not provide any additional insights on potential small molecule binding sites. Similarly, the 100 ns MD simulations do not show any evidence for large structural change. Thus, the well-characterized ETV6 PNT domain structure is retained both prior to and after polymerization.

### Alanine scanning mutagenesis identifies interfacial hot spot residues

Detailed SPR-monitored alanine scanning mutagenesis studies revealed that the ETV6 PNT domains heterodimerize with high affinity (K_D_ ∼ nM) due to residues partaking in electrostatic, van der Waals and hydrophobic interactions. Although it has been reported that hot spots are generally not enriched in electrostatic interactions (26), we found the opposite with residues in two proposed intermolecular salt bridges (K99-D101 and R105-D111) being critical for high affinity binding. Previous studies have shown that a K99R substitution also resulted in weaker binding, indicating that both the charge and amino acid structure are important, at least for one of these salt bridges (15). In contrast, the E100-R103 salt bridge does not contribute significantly as alanine substitutions of these residues did not result in a reduction in binding affinity or the oncogenic properties of the EN fusion protein (15). Indeed, presence of this salt bridge, and others involving these residues, varies between the reported ETV6 PNT domain structures. This also suggests that it is dynamic and not as persistent as the two formed by K99-D101 and R105-D111.

In addition to A93 and V112 (the founding sites for monomerizing mutations), several hydrophobic residues with the EH- and ML-surfaces are also hot spots for heterodimerization. In particular, L96 on the ML-surface plays a critical role in binding as removal of its side chain resulted in ∼ 1000-fold weaker affinity. The leucine makes contacts with several residues on the reciprocal EH-interface for which alanine substitutions also reduced the binding affinity, albeit each to a lesser extent. Targeting a molecule to bind near these EH-surface residues may exclude L96 and thereby inhibit PNT polymerization.

### Many of the hot spot residues have increased protection from amide HX

Alanine scanning mutagenesis and amide HX experiments provide complementary insights into the self-association of the ETV6 PNT domain. That is, alanine scanning revealed the effect of removing the side chain of a given residue on the affinity for heterodimer formation, whereas HX experiments showed how dimerization changes the conformational fluctuations of backbone amide hydrogens leading to exchange with water. In general, many of the hydrophobic hot spot residues within the core regions of the EH- and ML-surfaces, including L79, L96, L97, T98, Y104, V112, Y114, and L116 also exhibited enhanced HX protection upon heterodimerization. This is consistent with their burial, and likely dampened dynamics, within the heterodimer relative to the monomeric species. In contrast, although partaking in important intermolecular interactions, K99 and R105 underwent fast amide HX in both the monomeric and heterodimeric PNT domains, whereas their salt bridge partners D101 and D111 exhibited increased HX protection. This is also consistent with the location of the amides of these residues at the periphery of the dimer interface.

### Small molecule inhibition of PNT domain polymerization

Recently, we developed and implemented a mammalian cell-based assay, utilizing a protein-fragment complementation approach with split *Gaussia* luciferase, and a yeast two-hybrid assay in order to screen chemical libraries for potential inhibitors of ETV6 PNT domain self-association (27). In parallel, virtual screening using the Bristol University Docking Engine (BUDE) (28) was performed to identify compounds predicted to bind the ETV6 polymerization interfaces. Although numerous candidate molecules were tested, none had inhibitory effects in cellular assays and none bound to the isolated PNT domain. This work highlighted the difficulty in disrupting with small molecules the polymerization of the ETV6 PNT domain which occurs through two relatively large flat interfaces.

Insights into the factors that drive ETV6 PNT domain polymerization may aid inhibitor design. For example, small molecules that bind interaction surfaces near hot spot hydrophobic (e.g. L97) or salt-bridging (e.g. K99-D101 or R105-D111) residues may competitively prevent PNT domains from self-associating. However, designing or identifying such small molecules is challenging as the polymerization occurs with high affinity and there does not appear to be any structural differences between the PNT domains in their monomeric and heterodimeric states. One strategy that could be implemented for screening assays is to utilize an alanine mutant that weakens binding. This way, it may be easier to identify an initial compound that inhibits a micromolar affinity interaction, as opposed to the nanomolar interaction of the wild type PNT domains. Once discovered, such a lead molecule could be optimized for higher affinity binding. As previously shown, mutations of K99 weaken PNT domain polymerization and alter the oncogenic properties of EN (15). This residue is not at the center of the heterodimer interface and the K99A or K99R mutants may be well suited for such a screening strategy.

Alternatively, from the alanine scanning mutagenesis, we know which residues are not hot spots and thus tolerant to modifications. An example of a technique that would benefit from this acquired knowledge is disulphide tethering where weakly binding chemical fragments are tethered via an introduced cysteine residue near the protein-protein interaction interface (29). In principle, one could modify a residue, that does not affect binding and is near a hot spot, to a cysteine for identifying weak binders.

Helix “stapling” is another method that has been used to design molecules that disrupt protein-protein interactions. The general principle is to covalently stabilize the secondary structure of residues that normally form a helix along an interaction surface (30). The PNT domain, a subset of SAM domains, is a helical bundle, with helices H4 and helices H2 and H3 forming complementary interaction surfaces. Residues D111, V112 and Y114 in helix H4 are all hot spots and experience substantial increases in HX protection upon dimerization. Thus, a stapled helical polypeptide encompassing these residues might be sufficient to prevent polymerization. In contrast, whereas several residues on H2 and H3 are hot spots, such as D101 and R105, they are not adjoining as a single helix. A similar methodology has been used to target the Ship2 and EphA2 SAM-SAM domain interactions, whereby a penta-amino acid motif found in EphA2 binds to the SAM domain of Ship2, albeit with a K_D_ value in the high micromolar range (31). Therefore, exploration of polypeptide mimics of helix H4 for disruption of PNT domain polymerization may be a suitable option.

As a closing comment, SPR experiments demonstrated that the lifetime of the PNT domain heterodimer (1/k_off_) is ∼ 10 min. The lifetime of polymeric forms will be longer since dissociation must occur at multiple interfaces to completely monomerize. If the dissociation of the PNT domain polymer is slow in the context of a PNT-PTK fusion oncoprotein, then a prospective small molecule inhibitor that would have the greatest effect may likely need to act on newly translated PNT-PTK fusion oncoproteins, before they polymerize.

## Experimental Procedures

### PNT domain expression and purification

The DNA construct encoding residues 1-125 of human ETV6 (Genbank Gene ID: 2120), preceded by a thrombin-cleavable N-terminal His_6_ affinity tag, was initially cloned into the pET28a vector (Invitrogen) as described (15). The monomerizing A93D, V112E, and A93D-V112E substitutions were introduced through QuikChange site-directed mutagenesis (Stratagene). Subsequently, it was recognized that a thrombin cleavage site is present within the intrinsically disordered N-terminal region of these constructs (residues V37-P38-R39↓A40). This is located just before an alternative start site (M43) for ETV6 expression (11). Thus, ETV6 fragments were expressed from available clones as residues 1-125 with a His_6_ tag, and cleaved with thrombin to yield final purified samples spanning residues 40-125.

PNT domain-containing proteins were expressed in *E. coli* BL21 (λDE3) cells grown at 37 °C to an OD_600_ ∼ 0.6 and induced with 1 mM IPTG overnight. Unlabeled proteins were produced in lysogeny broth (LB) media, and M9 minimal media was supplemented with either 1 g/L of ^15^NH_4_Cl for ^15^N-labeled proteins or 1 g/L of ^15^NH_4_Cl and 3 g/L ^13^C_6_-glucose for ^13^C/^15^N-labeled proteins. In all cases, 35 mg/L kanamycin was included for plasmid selection. After continued growth at 37 °C overnight, the cells were harvested by centrifugation (Sorvall GSA rotor; 5,000 rpm) and frozen at −80 °C. The cell pellet was thawed for purification and resuspended in denaturing buffer (4 M GdnHCl, 20 mM sodium phosphate, 500 mM NaCl, 20 mM imidazole, pH 7.5) and lysed by sonication on ice. The lysate was then cleared by centrifugation (Sorvall SS34 rotor; 15,000 rpm) and the resulting supernatant was passed through either a 0.45 or 0.8 μm pore size filter and loaded onto a 5 mL Ni^+2^-NTA HisTrap HP column (GE Healthcare) pre-equilibrated with binding buffer (20 mM imidazole, 50 mM sodium phosphate, 500 mM NaCl, pH 7.5). After washing with several column volumes of binding buffer, the protein was eluted using a 120 mL linear gradient of elution buffer (500 mM imidazole, 50 mM sodium phosphate, 400 mM NaCl, pH 7.5).

The collected fractions were analyzed by SDS-PAGE and those containing the desired protein were pooled. In general, the ETV6 fragments eluted as two major peaks off the Ni^2+^ column. Previous studies demonstrated that proteins from these fractions had the same masses yet showed small differences in their ^15^N-HSQC spectra (32). Despite significant efforts, the origin of these differences was never elucidated. For consistency, the only fastest eluting peak was collected and dialyzed overnight at 4 °C in thrombin cleavage buffer (20 mM Tris, 0.15 mM NaCl, 2.5 mM CaCl_2_, 0.5 mM EDTA, 1 mM DTT, pH 8.4) with ∼ 1 unit of thrombin (Millipore) per 20 mL of collected fractions. The cleaved protein was separated from any His_6_-tagged species by passage through the Ni^+2^-NTA HisTrap HP column and purified further using size-exclusion chromatography (Superdex S75, GE Healthcare). This also served to exchange the protein in a buffer optimized for NMR experiments (noted below). The concentration of each protein sample was determined by ultraviolet absorbance at 280 nm based on its predicted molar absorptivity (33).

### General NMR spectroscopy methods

NMR spectra were recorded with cryoprobe-equipped Bruker Avance III 500, 600, and 850 MHz spectrometers. All data acquired were processed and analyzed with NMRPipe (34) and NMRFAM-Sparky (35, 36). Typically, protein samples were 150 μM to 600 μM in ∼ 450 μL of standard buffer (20 mM MOPS, 50 mM NaCl, and 0.5 mM EDTA) with D_2_O (5% v/v) added for signal locking. Unless otherwise noted, experiments involving the A93D-PNT domain and A93D-V112E-PNT domain were conducted at pH 7.0 and those involving the V112E-PNT domain were conducted at pH 8.0 due to its propensity to self-associate under lower sample pH conditions. In the case of the heterodimer species, the unlabeled partner protein was added to its isotopically-labeled partner in a 1.1 molar excess. The heterodimer containing an isotopically-labeled V112E-PNT domain was studied at pH 7.5 whereas that containing an isotopically-labeled A93D-PNT domain was studied at pH 7.0.

### Spectral assignments and chemical shift analyses

The signals from mainchain ^1^H, ^13^C, and ^15^N nuclei of the ^13^C/^15^N-labeled PNT domain constructs were assigned using standard heteronuclear ^1^H-^13^C-^15^N scalar correlation experiments (37), including HNCACB, HNCO, CBCA(CO)NH, HNCACO, along with HSQC-NOESY spectra, recorded on a 600 MHz spectrometer at 25 °C. Spectra of the monomeric A93D-PNT domain were assigned manually, whereas those of the monomeric V112E PNT domain and the two heterodimer complexes were automatically interpreted using PINE and verified manually (38). The assigned chemical shifts of these four species have been deposited in the BioMagResBank (BMRB) depository.

Secondary structure analyses were carried out using the MICS (Motif Identification from Chemical Shift) online server (17). Combined CSPs (Δδ) for the ^1^H^N^ (Δδ_H_) and ^15^N (Δδ_N_) signals of the corresponding amides in the dimer versus monomer species were calculated as Δδ = [(Δδ_H_)^2^ + (0.14Δδ_N_)^2^)]^1/2^.

### Amide hydrogen exchange (HX) by NMR spectroscopy

Protium-deuterium HX experiments for the PNT domains in the absence (monomeric) and presence of their unlabeled partner (heterodimeric) were conducted on a Bruker Avance 600 MHz spectrometer at 21 °C (matching ambient room temperature). The initial pH values were 7.0 for the samples containing the ^15^N-labeled A93D-PNT domains and 7.5 for those containing the ^15^N-labeled V112E-PNT domain. Initial reference ^15^N-HSQC spectra of the proteins in H_2_O buffer were recorded. Subsequently a 450 μL aliquot was lyophilized, resuspended with the same volume of D_2_O, and immediately put into the NMR spectrometer to begin data recording within 4-7 minutes. Initially, a series of ∼ 5-minute ^15^N-HSQC spectra were acquired with 2 scans/FID to characterize amides exchanging on the minutes timescale. Then ∼ 20-minute ^15^N-HSQC spectra with 8 scans/FID were collected back-to-back for ∼ 3 hours, followed by a 20-minute spectrum every hour for ∼ 24 hours, and then intermittent 20-minute spectra over a period up to 3 months. After the first week, the sample was removed from the spectrometer and stored at ambient room temperature between recording spectra. Upon completion of data recording, the pH* (pH meter reading uncorrected for the deuterium isotope effect) of each sample was measured as 7.3 (monomeric A93D-PNT domain), 7.6 (monomeric V112E-PNT domain), 7.4 (^15^N-labeled A93D-PNT domain in complex with V112E-PNT domain), and 7.5 (^15^N-labeled V112E-PNT domain in complex with A93D-PNT domain).

For each amide with measurable signals at times t after resuspension in D_2_O, the pseudo-first order exchange rate constant *k*_*obs*_ was obtained by fitting with NMRFAM-Sparky the ^1^H^N^-^15^N peak intensity *I*_*t*_, scaled for number of acquisitions/FID, to the equation for a single exponential decay:

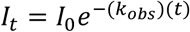

*I*_*0*_ is the fit initial intensity extrapolated to t = 0. The protection factor 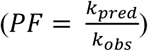 for each amide was determined as the ratio of its predicted intrinsic exchange rate constant (*k*_*pred*_) in an unstructured polypeptide of the same amino acid sequence versus its experimentally measured *k*_*obs*_. The *k*_*pred*_ values were determined with the program Sphere (39) which uses reference data based on poly-DL-alanine and corrected for amino acid type, temperature, pH and isotope effects (40, 41). In the cases of amide that had not exchanged significantly after 3 months, lower limits on their PFs were estimated based on the largest measured PFs for the given sample.

### Site-directed mutagenesis and construct cloning

To enable site-directed biotinylation during protein expression in *E. coli*, a gene encoding residues 43-135 of V112E ETV6 with an N-terminal His_6_-tag and Avitag was constructed in the pET28a vector using PIPE (polymerase incomplete primer extension) cloning techniques (42, 43). The A93D and E112V mutations were sequentially introduced to generate the complementary A93D-PNT domain construct with the His_6_-tag and Avitag.

Interfacial residues present in the X-ray crystal structure of the PNT domain dimer (PDB: 1LKY) were identified using the online SPPIDER (Solvent accessibility-based Protein-Protein Interface iDEntification and Recognition) server (44). Alanine substitutions were encoded at these sites in the respective A93D- or V112E-PNT domain clones using QuikChange site-directed mutagenesis techniques. All but two constructs were successfully generated in-house, and genes encoding N90A-V112E-PNT domain and L116A-A93D-PNT domain were purchased commercially (Biomatik).

### Protein expression and purification for surface plasmon resonance (SPR)

Each plasmid encoding either the biotinylated A93D-PNT and V112E-PNT domain was co-transformed into *E. coli* BL21 (λDE3) with the pET21a-BirA plasmid, which produces biotin ligase. Selection of both plasmids was maintained by including kanamycin (35 mg/L) and ampicillin (100 mg/L) in all media. Overnight seed cultures were used to inoculate LB media, supplemented with 0.05 mM biotin, and grown at 37 °C to an OD_600_ ∼ 0.6. Protein expression was induced with 0.5 mM IPTG and the cells were grown at 30 °C overnight. The cells were collected by centrifugation and cell pellets were frozen at − 80 °C.

Protein purification was carried out fully as described above with minor modifications. The final size exclusion purification step was omitted after thrombin cleavage and removal of the His_6_-tag by passage through a Ni^+2^-NTA HisTrap HP column. The final protein samples were exchanged into 20 mM MOPS, 50 mM NaCl, 0.5 mM EDTA buffer, at pH 8.0, and concentrated to ∼ 1 mL with an Amicon 3K MWCO centrifugal filter. The protein samples were then snap frozen in liquid nitrogen and stored at −80 °C prior to SPR analysis. The proteins were ∼ 50-95% biotinylated as judged by MALDI-ToF mass spectrometry.

### Surface plasmon resonance (SPR)

SPR experiments were performed on a Biacore X100 instrument using the streptavidin Sensor Chip SA to capture the biotinylated PNT domain “ligand”. The ligand was diluted to 50 μg/mL in HBS-EP+ buffer (10 mM HEPES, 150 mM NaCl, 3 mM EDTA, 0.05% v/v Tween-20, pH 7.4) and immobilized on the chip using the Biacore X100 control software immobilization wizard. The regeneration scouting software wizard was used to determine the regeneration step involving flowing 0.2% SDS over the chip for 30 s.

The association (k_on_) and dissociation (k_off_) rate constants and the equilibrium dissociation constant (K_D_) for binding of the analyte with the immobilized ligand were determined using the Biacore X100 kinetics/affinity assay software wizard. The positive and negative binding controls were analytes with the complementary wild type PNT domain interface or with the same monomerizing substitution, respectively. For the experimental runs, the analyte protein sample was initially diluted in HBS-EP+ buffer to 0.2, 2, 20, 40 and 60 nM. If weakened binding (K_D_ > 60 nM) was observed, the analyte protein sample was re-run in HBS-EP+ buffer at 20, 200, 2000, 4000 and 6000 nM.

### Structural determination of A93D-V112E-PNT domain by X-ray crystallography

A construct of ETV6 spanning residues 40-125 with the A93D and V112E substitutions (A93D-V112E-PNT domain) was purified as described above. Crystals were grown by sitting drop vapor diffusion at room temperature with 2 μL drops prepared with a 1:1 mixture of ∼ 9.3 mg/mL protein (20 mM MOPS, 50 mM NaCl, and 0.5 mM EDTA, pH 7.0) and reservoir solutions from the Hampton Index reagent crystallization screen (Hampton Research). Several conditions yielded protein crystals and those grown in 2.8 M sodium acetate (pH 7.0) were used for data collection. These crystals were cryoprotected by brief soaking in reservoir buffer supplemented with 30% (v/v) glycerol, followed by flash freezing in liquid nitrogen.

Diffraction data were collected at the CLS (Canadian Light Source) on beamline 08B1-1 (45). The data were cut-off at 1.85 Å based on the CC1/2 metric and processed and scaled using XDS (46). Crystals were of space group *P*6_5_22 with two protein molecules in the asymmetric unit. Phase determination using molecular replacement was performed with a PNT domain monomer from PDB: 1LKY using the AutoSol program in Phenix (47). Model building was performed in Coot and refinement was executed using the Phenix software suite (48). Eight of 85 amino acid residues of the construct (4 at the N-terminus and 2 at the C-terminus) were disordered in the crystal and not modelled.

### Molecular dynamics (MD) simulations on monomeric and dimeric PNT domains

MD simulations were based on coordinate files (that encompassed from residues S47 to Q123) from the following crystal structures: the monomeric A93D-V112E-PNT domain (determined herein), an A93D-PNT domain subunit from PDB: 1LKY (with an alanine introduced at position 93 with the mutagenesis function in PyMol (Version 1.8) (49)), and a dimer of the A93D-PNT domain and V112R-PNT domain from PDB: 1LKY.

All MD simulations were performed on the University of Bristol High Performance Computer BlueCrystal using GROMACS (Version 5.1.2) (50). The systems were solvated with TIP3P waters in an orthorhombic box 2 nm larger than the longest dimension of the protein. Sodium and chloride ions were included at 50 mM to emulate experimental conditions, while neutralizing the monomerizing mutations to have no overall net charge. The amber99sb-ildn forcefield was used to parameterize the protein simulations (51). Non-bonded long-range electrostatic interactions were calculated using the Particle Mesh Ewald method with a 1.2 nm cut-off. Bonds were constrained using the LINCS default algorithm to allow the use of a 2 fs timestep for the MD integration.

The energy of the system was minimized over 5000 steps of the steepest descent energy minimization. The system then underwent a position-restraint simulation over 200 ps where the protein was restrained to its initial position while heating the system to 310 K and introducing pressure at 1 bar using the Berendsen barostat (50). The full unconstrained MD simulations were run over 100 ns with integration step sizes of 2 fs using the leap-frog algorithm, and trajectory files were recorded every 100 ps. The temperature was maintained at 310 K using v-rescale modified Berendsen thermostat temperature coupling and at 1 bar with the Berendsen barostat pressure coupling. Trajectories were analyzed and processed, including principal component analysis, utilizing GROMACS tools. The simulations were visualized with VMD (Version 1.9.2) (52), gnuplot (Version 4.6) and PyMol (Version 1.7) (49).

### Data Availability

The X-ray structure (coordinates and structure factor files) has been submitted to the PDB under accession number 7JU2. The NMR chemical shifts have been submitted to the BMRB under accession numbers 50430 (Monomeric V112E-PNT domain), 50431 (Complexed V112E-PNT domain), 50432 (Complexed A93D-PNT domain), and 50433 (Monomeric A93D-PNT domain). All other data described here are available within the manuscript and supporting information.

## Acknowledgements

We thank Dr. Poul Sorensen (British Columbia Cancer Research Centre) for advice on ETV6 translocations, and the Advanced Computing Research Centre at Bristol University for the provision of high performance computing.

Supporting information is available online with this article.

## Author Contributions

C.A.N.G., L.P.M., and M.R. were responsible for the conceptualization, data analysis, supervision, and writing of this manuscript. C.A.N.G. and M.O. carried out the NMR experiments. S.Y.C. and C.A.N.G., preformed the alanine scanning mutagenesis experiments, and C.A.N.G. and M.K. undertook the x-ray crystallographic experiments, under guidance of M.E.P.M. The MD simulations were run by C.A.N.G. with assistance from R.B.S.

## Funding

The authors disclosed receipt of the following financial support for the research, authorship, and/or publication of this article: This study was supported by funds from the Canadian Cancer Society Research Institute to L.P.M and M.R., and from the Canadian Institutes of Health Research (CIHR) to L.P.M. C.A.N.G. held graduate scholarships from CIHR and the University of British Columbia.

## Declaration of Conflicting Interests

The authors declared no potential conflicts of interest with respect to the research, authorship, and/or publication of this article.

## Notes

### Competing Interest Statement

The authors have declared no competing interest.

